# An Analytic framework for stochastic binary biological switches

**DOI:** 10.1101/068759

**Authors:** Guilherme C.P. Innocentini, Sarah Guiziou, Jerome Bonnet, Ovidiu Radulescu

## Abstract

We propose an analytic solution for the stochastic dynamics of a binary biological switch, defined as a DNA unit with two mutually exclusive configurations, each one triggering the expression of a different gene. Such a device could be used as a memory unit for biological computing systems designed to operate in noisy environments. We discuss a recent implementation of an exclusive switch in living cells, the recombinase addressable data (RAD) module. In order to understand the behavior of a RAD module we compute the exact time dependent distributions of the two expressed genes starting in one state and evolving to another asymptotic state. We consider two operating regimes of the RAD module: fast and slow stochastic switching. The fast regime is “aggregative” and produces unimodal distributions, whereas the slow regime is “separative” and produces bimodal distributions. Both regimes can serve to prepare pure memory states when all cells are expressing the same gene. The slow regime can also separate mixed states by producing two sub-populations each one expressing a different gene. Our model provides a simplified, general phenomenological framework for studying biological memory devices and our analytic solution can be further used to clarify theoretical concepts in bio-computation and for optimal design in synthetic biology.

---

Biological machines are a promising new paradigm in computation [11]. By using synthetic biology it is now possible to design memory units and logic gates operating in living cells [6, 7]. Contrarily to their silicon counterparts, biological computing units (BCU) can evolve their computation capacities by proliferation and auto-organization. Furthermore, parallel computing with large number of BCU could be used to performing distributed computation in living environments for medical applications such as implants for augmenting capacities or for health monitoring [4].

However, there are prices to pay when replacing the silicon substrate with biological cells. BCUs are submitted to stochastic fluctuations that are ubiquitous in the realm of molecular machines [8], [10]. Within cell communities, the result of a computation can vary from one cell to another and therefore this result should be represented as a probability distribution between the different states of the system. In order to optimize the design of BCUs, precise calculations of time dependent population distributions is extremely useful. In this paper we show how to solve this problem in the context of the set and reset of a reversible and stochastic, rewritable biological unit.

Synthetic biological memory devices have been engineered based on different mechanisms. Some systems use feedback loop in order to achieve data storage via bistability. For example, double negative feedback systems inspired by the phage lambda Cro/CI circuit and using two repressor mutually repressing each others have been implemented in the bacterium *Escherichia coli* [9], [12]. Positive feedback based memory devices have also been engineered in the baker’s yeast *Sacharomyces cerevisiae* [2]. Recently, genetic memory systems using recombinases from the integrase family, enzymes that catalyze strand exchange between specific DNA sequences [4], have been implemented into living cells. DNA data storage has the advantage that only two discrete states are possible. The state of the system can be interrogated by direct DNA sequencing but also by measuring the intensity of a fluorescent protein (FP) whose expression is controlled by the invertible DNA sequence (called the DNA register). Fluorescent reporters of two colors, for instance Green Fluorescent Protein (GFP) and Red Fluorescent Protein (RFP), can be used to signify binary DNA register states of single cells. Although the majority of DNA data storage devices are single-write units, a particular architecture called the recombinase addressable data (RAD) module using two proteins, an integrase and a recombination directionality factor (or excisionase) enables rewritable digital data storage in bacteria [5] (Fig.1).

**FIG. 1:**
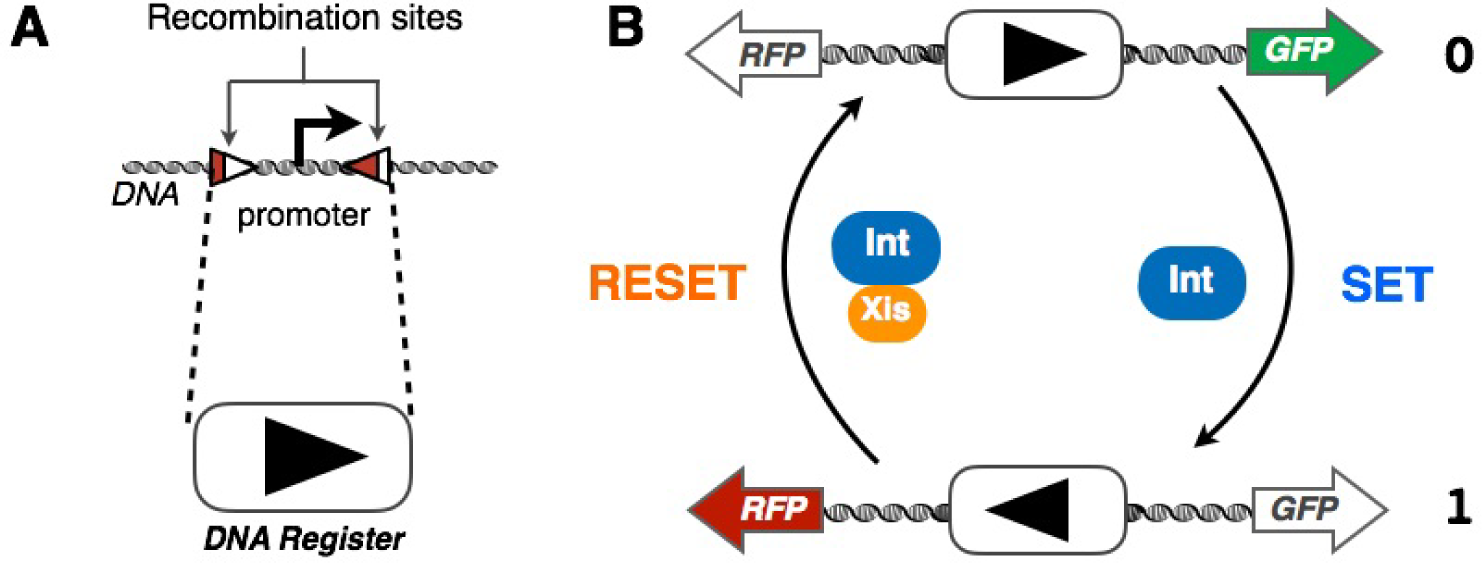
Schematic architecture and operational principle of the Recombinase Addressable Data (RAD) module. A. The DNA register is composed of a DNA sequence containing a promoter (driving transcription) flanked by the two recombination sites recognized by the integrase (Int). B. In state 0, the promoter drives expression of GFP. Integrase catalyzes the SET reaction, inverting the DNA register and enables transition towards state 1, in which the promoter has been inverted and now drives transcription of RFP. Concomitant expression of Integrase and Excisionase (Xis) catalyze the RESET reaction by enabling transition from state 1 to state 0.

The RAD module proposed by Bonnet et al [5] is composed of a DNA sequence which can exist in two configurations inverted one with respect to the other. The transition between the configurations is reversible. Depending on the configuration, single cells produce one of the two reporters, a GFP or RFP (Fig.1). Although the two internal states of the memory defined by the DNA configurations are mutually exclusive, the expression of the two reporter genes depends on how fast the unit switches between the two configurations. As we will prove later in this letter, fast switching between the two configurations allow that both GFP and RFP are expressed by the same cell. In this work, we are interested in the conditions enabling the generation of cell populations residing in a pure state (when all the cells express the same reporter gene) or separated mixed state (when sub-populations express an unique reporter gene but not necessarily the same). To this end, we introduce a mathematical model to describe the stochastic dynamics of the system. Our model is based on two coupled master equations governing the evolution of the joint probability distributions carrying the information about switch status and numbers of GFP (*g*) and RFP (*r*) at a given instant of time:

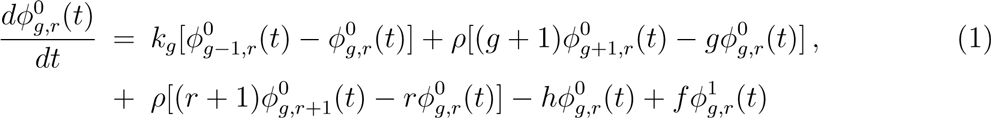

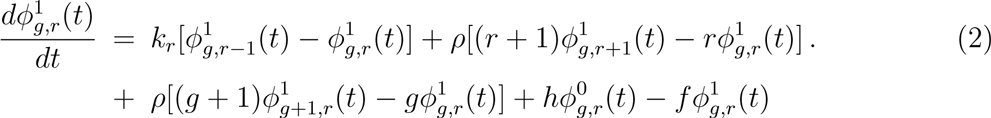

The random variables are *g* and *r*. Production of GFP and RFP is controlled by the rates *k*_*g*_ and *k*_*r*_, respectively. Degradation/dilution rate of both reporters is given by *ρ* and the switching between the two states is encoded in the rates *h* (SET) and *f* (RESET).

Introducing the generating functions, 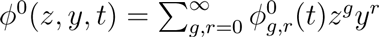 and 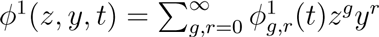, we transform the master equations in a set of partial differential equations:

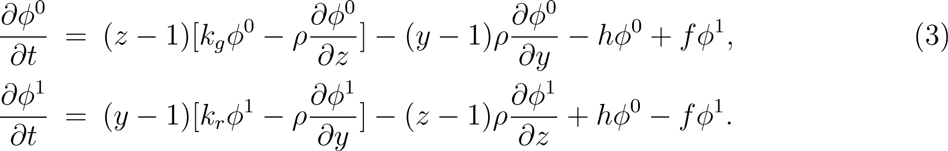

The desired joint probability distributions are obtained from the generating functions by applying the formulas:

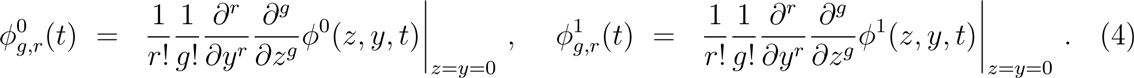

In order to solve the system of PDEs in Eq.(3) we propose a set of transformations, that will lead to an integrable system of ODEs. To do so, we perform a first change of variables:

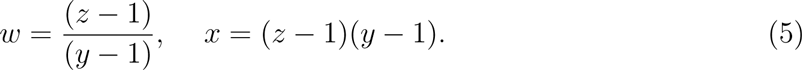

In the new set of variables (*w, x*) the equations read:

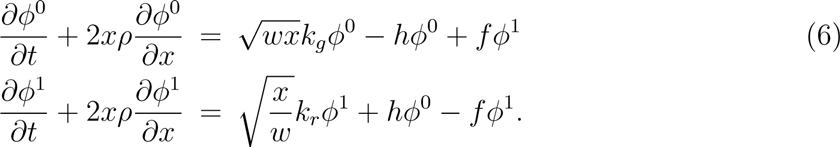

Note that this transformation let time (*t*) unchangeable and had the effect of eliminating one of the partial derivatives. To eliminate another partial derivative and obtain a set of ordinary differential equation with respect to the remaining variable we propose a second transformation,

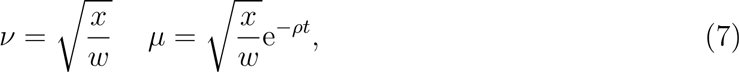

which leads to the set of ordinary differential equations in the variable *v*:

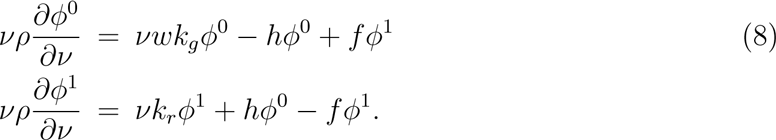

Now, solving the first equation of the system (8) for *ϕ*^1^(*ν, μ, w*), gives

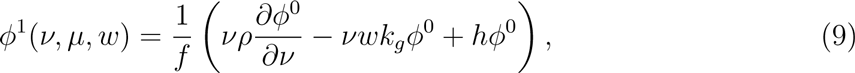

and by substituting the result in the second equation of the system, we arrive to the second order ordinary differential equation with respect to the variable *ν* for *ϕ*^0^(*ν, μ, w*):

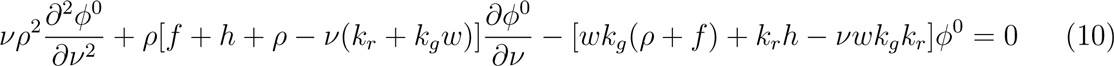

This equations has a regular singularity at *v* = 0 and a irregular one at infinity. This structure suggest solutions in terms of confluent hypergeometric functions. To make it more clear, let us use the ansatz *ϕ*^0^(*ν, μ, w*) = exp(*νwk_g_*/*ρ*)*ψ*(*ν, μ, w*), and a last transformation of variables: *ν = ηρ*/(*k_r_ – wk_g_*). Putting all together, we arrive to the equation in the new variable *η* for *ψ*(*η, μ, w*):

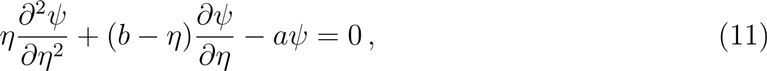

where: *a = h*/*ρ, b* = (*h + f + ρ*)/*ρ* and *η* = *ν*(*k_r_ – wk_g_*)/*ρ*.

Now, Eq.(11) is in the canonical form of the confluent hypergeometric equation, or Kummer equation, and the general solution is straightforward:

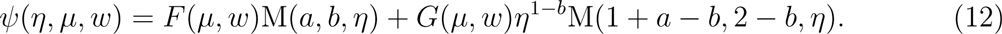

where M stands for Kummer function, *F*(*μ, w*) and *G*(*μ, w*) are arbitrary functions that will be determined by the initial conditions.

To obtain the expressions for the generating functions we have to multiply the solution in Eq.(12) by the exponential factor, exp(*νwk_g_*/*ρ*), to obtain *ϕ*^0^(*η,μ,w*) using this result and Eq.(9) we can obtain the generating function for *ϕ*^1^(*η,μ,w*), in the variables (*η,μ,w*):

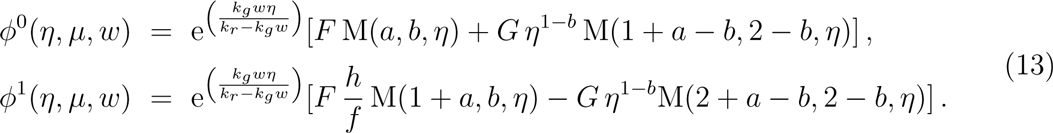

The last task remaining is to find *F*(*μ,w*) and *G*(*μ,w*). As said before, these functions are determined by the initial conditions. To accomplish that, one can see that setting *t* = 0 in the second transformation Eq.(7) leads to *v = μ*, which in the variables (*η,μ,w*) implies in 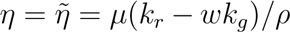. Specifying the functions 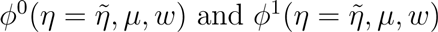 that will appear in the left hand side of Eq.(13) one can find the expressions for *F*(*μ,w*) and *G*(*μ,w*). To do so, let us use vector and matrix notation to express the solutions in Eq.(13) as: 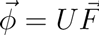, where, 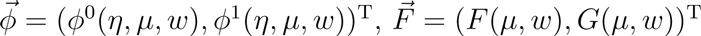 and the entries of the matrix *U* are given by:

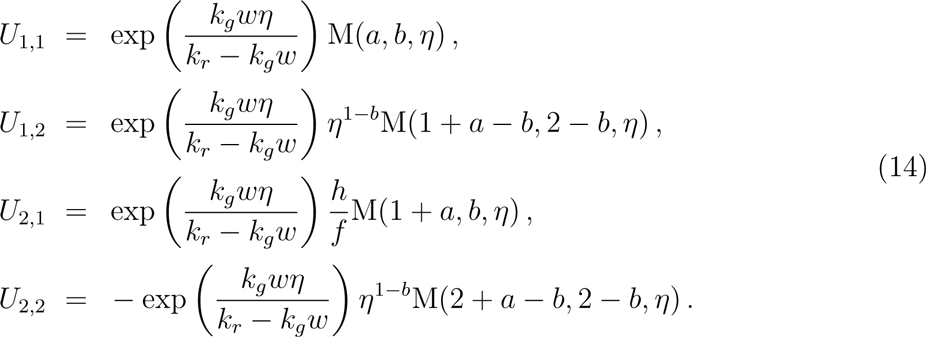

Inverting 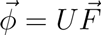 we obtain the expression 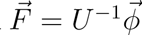. Setting 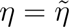 bring us to the position of determining the vector 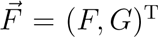 through the initial conditions 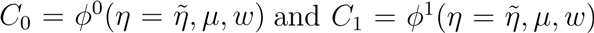. One final observation, before presenting *F* and *G*, concerns the determinant of the matrix *U* which is necessary because we need this determinant (det(*U*) = *U*_1,1_*U*_2,2_ — *U*_1,2_*U*_2,1_) to compute *U*^−1^ (the inverse matrix of *U*). The inspection of (14) reveals that *det*(*U*) is a product of Kummer functions with a exponential envelope. At a first glance, no problem arise, however, as we will deal with the inverse of matrix *U* the inverse of this determinant is required. Due to the well known properties of the Kummer functions [1] this determinant assumes a very simple formula:

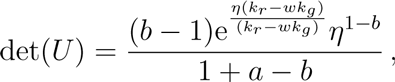

which is used to compute *U*^−1^. The explicitly expressions for *F* and *G* are:

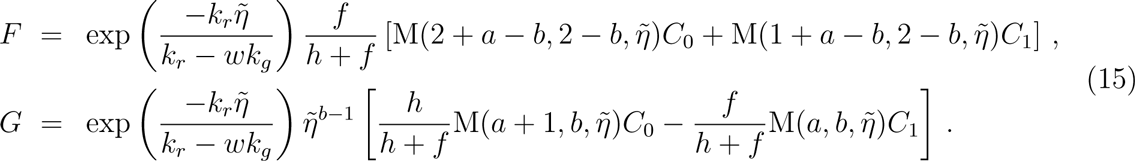

Where initial conditions are encoded in 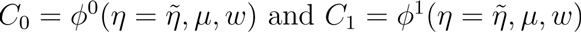.

At this point we are in position to exhibit the biological features of our model encoded in the time dependent joint distributions of three variables: the two-valued DNA register state and the numbers of GFP (*g*) and RFP (*r*). However, before doing that, let us rephrase our parameter space in adimensional biological terms. The two numbers *N_g_* = *k_g_/ρ* and *N_r_* = *k_r_/ρ* are the protein production efficiencies. The asymptotic occupancy probabilities are *p_0_* = *f*/(*f* + *h*) and *p*_1_ = *h*/(*f + h*). We call *switching flexibility* the important parameter ∈ = (*h + f*)/ρ, representing the sum of the frequencies of the SET and RESET transitions. The two situations ∈ < 1 and ∈ > 1 correspond to slow and fast switching, respectively.

Although we have the time dependent solutions at hand we will first illustrate the role of the biological parameters by analyzing their influence in the shape of the asymptotic distributions of the model. The generating function for the the steady-state distributions is obtained by performing the limit *t* → ∞ in Eqs.(13), resulting in:

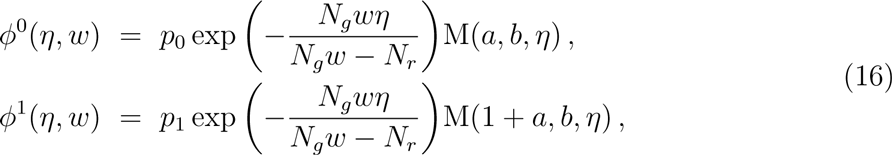

where, in the biological parameter space we have: *a* = *p*_1_*ϵ b* = *ϵ* + 1, and *η* = *ν*(*N_r_ – wN_g_*). Applying formulas (4) we obtain the steady-state joint probability distributions as

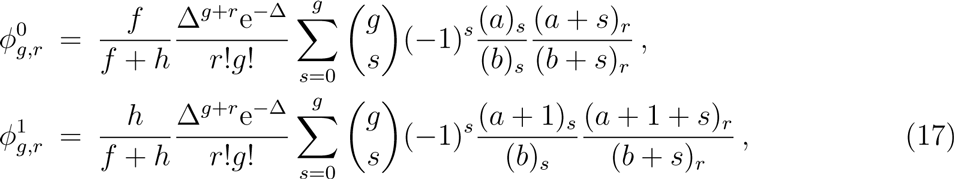

where we have used *N_g_* = *N_r_* = Δ and the symbol (•)_*s*_ is the Pochhammer’s symbol [1].

With Eqs.(17), we have computed the total joint probability distribution in the asymptotic regime of the system for different values of *p*_0_, *p*_1_ and ∈ keeping Δ constant. The steady-state distributions are represented in Figs.2a-c. In order to expand the regions of small expression we made a change of variables in the distributions from *g, r* variables to log(*g*), log(*r*) variables. As said before, we are doing this exercise to show the influence of the parameters in the shape of the distributions and, moreover, to distinguish between mixed and pure states of the DNA register. This concept is important for the relation between hidden memory states and visible (readable) phenotype. Let us define a mixed state by 0 < *p*_0_ < 1 and 0 < *p*_1_ < 1, whereas a pure state means that either *p*_0_ = 1 or *p*_1_ = 1. Also, we define three types of sub-populations regarding the level of expression of: GFP only; RFP only and both GFP and RFP. The corresponding state occupancies are *p_g_*, *p_r_*, *p_rg_*, respectively. A well separated population has low values of *p_rg_*. The state occupancies are not necessarily the same as the occupancy probabilities *p*_0_ and *p*_1_ encoding the probability to find the DNA register in state 0 or 1, respectively. A RAD unit in a pure state has a pure phenotype (*p_g_* = 1 or *p_r_* = 1), in other words all the bacteria express GFP in state 0 or RFP in state 1, as show in Fig.2a,b. The readout of a mixed state will depend on the switch flexibility. A fast switch (∈ > 1) will correspond to unimodal distribution of expressed genes, Fig.2c (each bacteria expressing both genes in different proportions) whereas a slow switch (∈ < 1) corresponds to bimodal population, Fig.2d, where some cells are green and others are red.

**FIG. 2:**
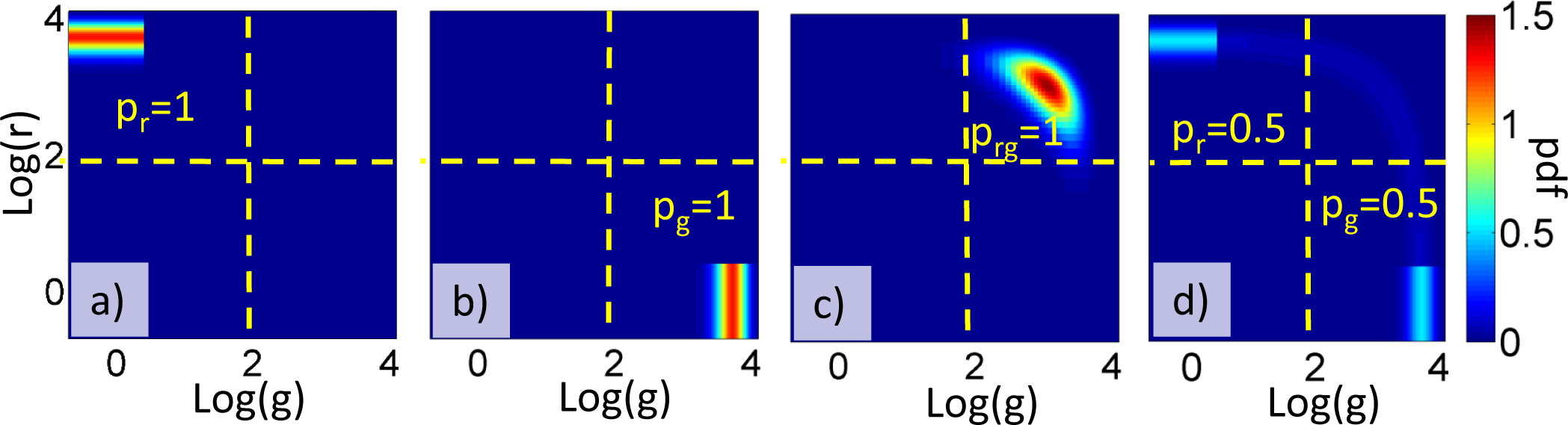
Computed steady state distributions of GFP and RFP for a) pure *p*_1_ = 1 red state (all cells express RFP) and ∈ = 0.5 (slow switching); b) pure *p*_1_ = 1 green state (all cells express GFP) ∈ = 0.5 (slow switching); c) mixed *p_0_* = *p*_1_ = 0.5 unimodal state (all cells express both GFP and RFP) and ∈ = 10 (fast switching); d) mixed p_0_ = p_1_ = 0.5 bimodal state (half of the cells express only GFP and half only RFP) and ∈ = 0.5 (slow switching). In logarithmic variables, the double integral of the probability distribution is normalized to one. The remaining parameters are: Δ = 40 and *ρ* = 1. The probabilities *p_r_*, *p_g_* and *p_rg_* indicate the fractions of cells with *log*(*g*) < 2,*log*(*r*) *>* 2, *log*(*g*) > 2,*log*(*r*) < 2 and *log*(*r*) > 2,*log*(*g*) > 2, respectively, where the threshold 2 corresponds to the middle of the expression dynamical interval.

In order to illustrate the dynamical behavior of the RAD module we will first set the initial conditions (*C*_0_ and *C*_1_) of the system. To do so, we use the steady-state solutions as initial conditions but with different values for the occupancy probabilities and switch flexibility 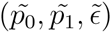, leading to the initial generating functions:

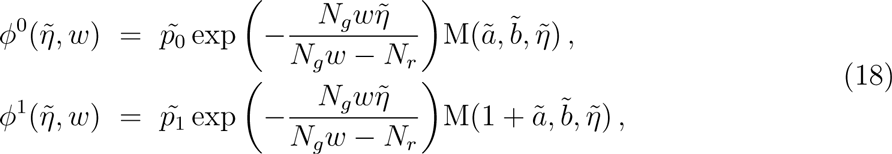

At *t* = 0 we abruptly change the occupancy probabilities and switch flexibility from 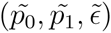 to (*p*_0_, *p*_1_, *ϵ*), keeping them constant for *t* ≥ 0. To study set and reset dynamics of the RAD unit we consider three distinct experiments and we show the corresponding time dependent joint probability distributions for each one of these experiments in Fig.3. The first two experiments correspond to setting a pure state. We start with initial condition corresponding to a pure state 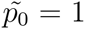 where all cells are producing GFP (in the setting described in Fig.1 this means that a strong excisionase signal is applied together with the integrase), and at time t = 0 we change the asymptotic occupancy parameter to the complementary pure state, *p*_1_ = 1, driving all the cells to be producing RFP in the asymptotic configuration (excisionase is washed-out). During the setting we use fast switching (*ϵ* = 10) for the first experiment Fig.3(a) and slow switching (*ϵ* = 0.5) for the second one Fig.3(b) (corresponding to high and low concentrations of integrase). In the third experiment, Fig.3(c), we start with an unimodal mixed steady-state configuration, in the fast switch regime (*ϵ* = 10, corresponding to high integrase and excisionase concentrations) and change the switch flexibility to low values (*ϵ* = 0.1, lower concentrations). We call this last experiment “developing” because it transforms the initial unimodal mixed state in which single cells express both GFP and RFP (state occupancy *p_rg_* = 1) into a bimodal, “separated” mixed state when two subpopulations have pure phenotypes, expressing either RFP or GFP (*p_r_* = *p_g_* ≈ 0.5 in Fig.2). More generally, the final state occupancies are given by the DNA occupancy probabilities *p_r_* ≈ *p*_1_,*p_g_* ≈ *p*_0_. Therefore, lowering the switching frequencies by lowering the integrase and excisionase concentrations reveals previously hidden information about the DNA register occupancy probabilities. The last column of Fig.3 shows the time dependence of the state occupancies (*p_g_* (*t*), *p_r_*(*t*) and *p_rg_* (*t*)) corresponding to each one of the three experiments.

**FIG. 3:**
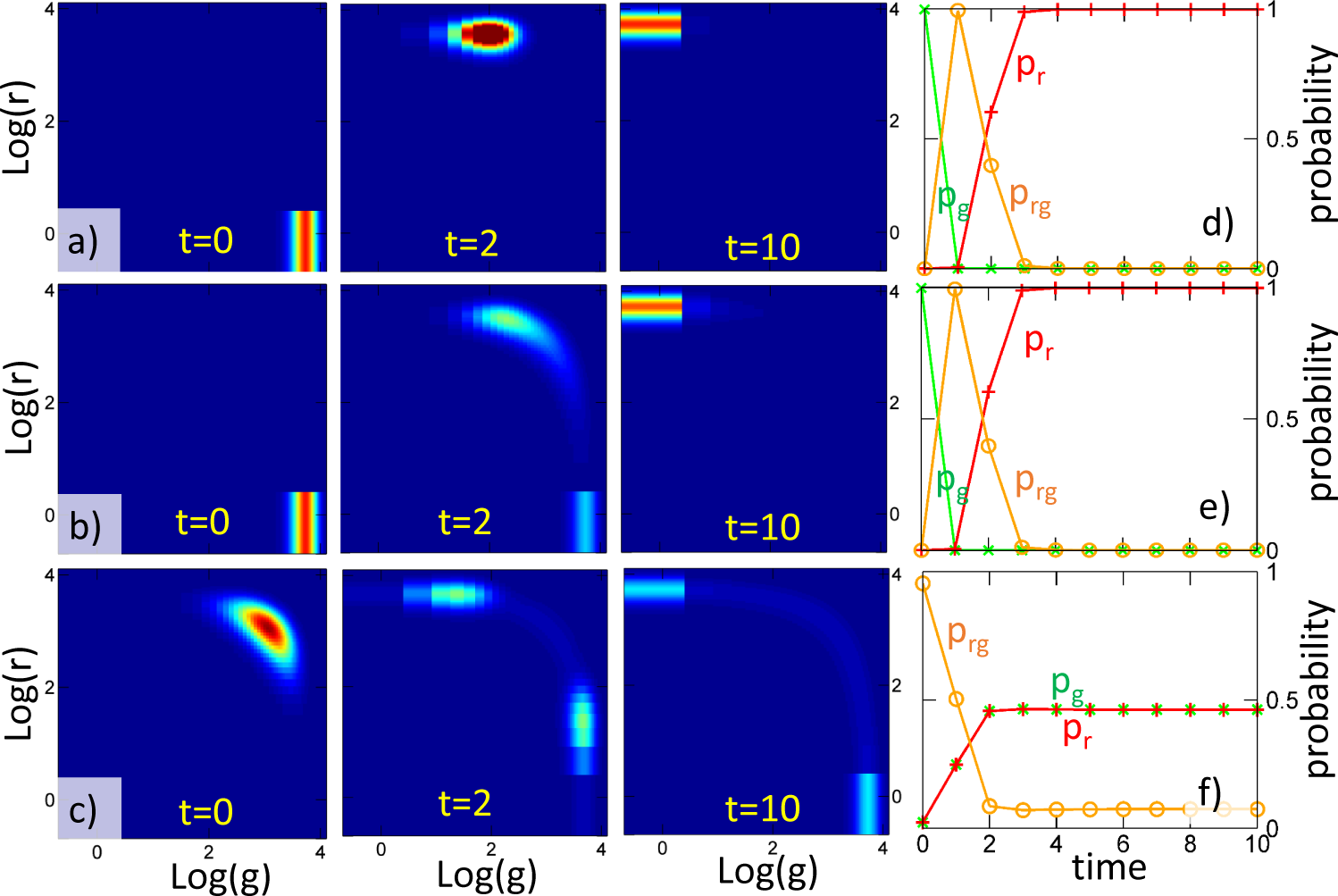
Time dependent distributions of a RAD unit (time is measured in units of *ρ*^−1^). In a),b) we illustrate setting of a pure state. The *t* = 0 state corresponds to a pure distribution with 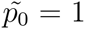 (green) and switch flexibility 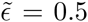. During set operation (for *t* > 0) the unit evolves towards a pure state *p*_1_ = 1 (red) but with different switch parameters: *ϵ* = 10 for a) (fast switching) and *ϵ* = 0.5 for b) (slow switching). In c) we illustrate separation of a unimodal mixed state. An unimodal mixed state with 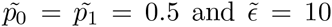 (fast switching) is transformed to a bimodal mixed state by lowering the switch flexibility, *ϵ* = 0.1 (slow switching). Also, in c) one can see that for fast switching the phenotype is unimodal: bacteria express both GFP and RFP. The *t* = 10 phenotype, is bimodal: 50% of the cells express RFP and 50% express GFP. The remaining parameters are: *N_g_* = *N_r_* = 40 and *ρ* =1. The rightmost columns d-f) show the time dependence of the state occupancies.

These experiments emphasize a clear distinction between slow and fast switches. Slow switches are “separative”, they tend to transform unimodal distributions into bimodal ones. They can be used as developers of mixed states, as occupancy probability can be read on bimodal distributions by counting cells in the two sub-populations, whereas it is much more difficult to estimate it from unimodal distributions. This can also be seen from the values of the state occupancies *p_rg_*(*t*) that remain low (indicating separated population) during the dynamics. For slow switches there is a trade-off between good separation and bandwidth because the time needed to reach a final unimodal, pure state distributions is longer. Fast switches are “aggregative”, they tend to transform bimodal distributions into unimodal distributions.

The analytic time dependent solutions for the master equations describing the RAD memory unit can be used to quantitative and qualitatively establish the basis for the design of such biological devices. All the necessary information about the system is encoded in the time dependent joint probability distributions which are obtained by applying formulas (4) in the analytic solutions of the model. The solutions are expressed in closed form in terms of the well known Kummer functions and packages in softwares of symbolic computation, such as Maple, are available for direct computation of their series expansions leading to the desired joint distributions. If, instead of evaluating the derivatives of the solutions in *z* = *y* = 0 as in formulas (4), one evaluates them at *z* = *y* = 1, closed expressions for any desired moment of the joint distributions, such as mean values, variance and covariance are easily obtained. A special word should be said about “developing” of a mixed state using slow switching. Although this procedure has no direct interest for storing information, it could be used as a biological implementation of a p-switch. A p-switch [13] is a stochastic element that takes two values, one and zero with probabilities *p* and 1 − *p*, respectively. In other words, it is a physical implementation of a Bernoulli random variable, which is the basic element in stochastic computing [3], an hybrid, digital/analog form of computation in which real numbers are represented as random finite sequences of zeros and ones, the represented real number *p* being the frequency of occurrence of ones. In our case, each random bit in a sequence would be a cell. Although redundant from an information theoretic point of view, this representation was shown to be particularly robust and protected against random faults [3].

Although the discussion is based on the RAD switch setting illustrated in Fig.1, our model provides a simplified phenomenological framework for studying other implementations of two states memories, including toggle-switches based on positive feedback. In the latter case, simplified switching models corresponding to stochastic transitions between the two attractors of the system can be justified by low noise approximations as discussed in [10].

## Acknowledgements

Work supported by FAPESP, SP, Brazil (G.I., contract 2012/04723-4) and CNPq, Brazil (G.I., contract 202238/2014-8). O.R. thanks LABEX Epigenmed for support.

